# Human alpha-defensin 5 stabilizes the enterovirus A71 capsid and blocks infection

**DOI:** 10.64898/2026.07.02.735982

**Authors:** Kaitlin R. Hulce, Manasa Acharya, Jessica M. Porter, Jason G. Smith

## Abstract

Human alpha-defensins are antimicrobial peptides abundantly expressed in neutrophils and the small intestine. They block infection of several families of non-enveloped DNA viruses by binding to and stabilizing the viral capsid during entry, thereby preventing the genome from reaching the nucleus to initiate replication. It is unclear if a similar mechanism also applies to RNA viruses. To study this further, we investigated the interaction of human alpha-defensin 5 (HD5) with enterovirus A71 (EV-A71). We found that HD5 disrupts EV-A71 infection in cell culture and blocks viral entry. HD5 binds directly to the EV-A71 capsid and disrupts key conformational changes essential to the initiation of *in vitro* uncoating as well as downstream viral genome release. Using a suite of HD5 point mutants, we found that these two uncoating blocks are separable, and HD5 must achieve both to fully neutralize EV-A71 infection. This work advances our understanding of alpha-defensin antiviral action and demonstrates several conserved features of HD5 inhibition that expand to a clinically important RNA virus.

**Author Summary:** In this study, we demonstrated that human defensin 5 (HD5) inhibits infection of pathogenic enterovirus A71 (EV-A71). HD5 binds to EV-A71 and disrupts *in vitro* viral uncoating by stabilizing the viral capsid and blocking genome release. Using a library of HD5 mutants, we identified a continuous surface within HD5 important for binding to EV-A71 and disrupting infection. This surface includes amino acid residues previously shown to be important for inhibition of human adenovirus 5 and human papillomavirus 16 and represents a conserved interface involved in broad-spectrum HD5 antiviral activity. We determined that capsid stabilization and blocked genome release are independent modes of HD5 action and identified individual HD5 mutants with separable functions. These mutants provide valuable tools to further study the biology of EV-A71 uncoating. Overall, this work demonstrates the first example of an enterovirus inhibited by an α-defensin and advances our mechanistic understanding of HD5 antiviral action to a new pathogenic virus.

## Introduction

Defensins are a ubiquitous family of antimicrobial peptides (1,2). In vertebrates, there are three subfamilies of defensins (α-, β- and θ-defensins), which are small, cationic, folded peptides of 18-45 amino acids in length (1–3). The antimicrobial activity across defensin subfamilies is complementary but not fully overlapping. While all three subfamilies neutralize bacterial and enveloped viral infection, evidence for activity against non-enveloped viruses is largely restricted to α-defensins (2–6). There are six human α-defensins, which can be further classified as either myeloid or enteric (3–5). Myeloid α-defensins, including human neutrophil peptides 1 to 4 (HNP1 to HNP4), are primarily expressed in neutrophils and secreted in response to pathogens (4). Enteric α-defensins, including human defensin 5 and 6 (HD5 and HD6), are constitutively expressed and secreted by Paneth cells into the crypts of the small intestine and can also be found in the genitourinary tract (5). Physiological concentrations of α-defensins range from high nanomolar (plasma) to low millimolar (phagolysosomes), with observed *in vitro* antimicrobial potencies falling within this range (3–5,7).

The neutralizing activity of α-defensins against non-enveloped viruses involves direct protein-protein interactions with the viral capsid, often leading to capsid stabilization (7–17). This mechanism is distinct from their activity against bacteria and enveloped viruses, which involves interactions with membrane lipids or glycans (2,4–6). While α-defensin activity has been characterized for several representative DNA viruses from the *Adenoviridae*, *Papillomaviridae*, *Parvoviridae*, and *Polyomaviridae* families, studies of RNA viruses are limited. We have recently demonstrated that α-defensins neutralize rotaviruses, but the mechanism remains unclear (18). To better understand the effects of α-defensins on RNA viruses, here we expand our studies to include enteroviruses (EVs).

EVs are a large, diverse group of positive sense RNA viruses belonging to the *Enterovirus* genus of the *Picornaviridae* family (19). The genus consists of fifteen species, seven of which cause diseases in humans (20). These include enteroviruses, coxsackieviruses, echoviruses, and polioviruses (species EV-A through EV-D), as well as rhinoviruses (species RV-A through RV-C) (21). EV infection in young children and immunocompromised individuals can lead to severe complications including acute respiratory diseases, serious neurological conditions, and heart complications (19). Of particular concern is enterovirus A71 (EV-A71), which is one of the etiological agents of hand-foot-and-mouth disease and has caused several serious outbreaks across the Asia-Pacific region (20,22,23). EV-A71 is highly neurotropic, and infection can lead to aseptic meningitis, acute flaccid paralysis, and encephalitis (24,25). Furthermore, one of the major causes of EV-A71 related mortality in young children is viral-induced pulmonary edema and hemorrhage (25). Though there are three EV-A71 vaccines approved in China, there are currently no FDA-approved vaccines in the US nor specific antivirals to manage infection (26,27).

Only one published study has examined the effects of α-defensins on EV infection, demonstrating that echovirus 11 is resistant to HNP1 (28). To expand our understanding of EV/α-defensin biology, we investigated EV-A71 because of its clinical relevance. We found that HD5, but not HNP1, has neutralizing activity against EV-A71 infection and blocks viral entry. We determined that HD5 binds to and stabilizes the EV-A71 capsid, leading to disrupted viral uncoating *in vitro* by blocking both heat-triggered particle expansion and genome release. Using point mutations of HD5, we found that these functions are separable and that both are necessary for complete inhibition of EV-A71 infection. We therefore posit a mechanism of action whereby HD5 binds to the EV-A71 capsid and blocks uncoating, leading to disrupted infection. This work suggests that the mechanism of action observed against DNA viruses also extends to ΕV-A71.

## Results

### HD5 neutralizes EV-A71 but not EV-D111 infection

We selected HNP1 and HD5 as representative human myeloid and enteric α-defensins, respectively, and tested their dose-dependent effects on EV-A71 infection of human rhabdomyosarcoma (RD) cells. HNP1 is abundantly expressed in neutrophils, while HD5 is one of the two human enteric α-defensins and is a more potent antiviral (5). We tested the effects of HD5 and HNP1 on EV-A71 infection between physiologically relevant concentrations of 1 to 40 μM (3–5,7). HNP1 caused a modest, biphasic effect on EV-A71 infection, with a 2-fold increase between 1 and 10 μM and a 50% reduction at 40 μM (**Fig 1A**), indicating overall weak neutralizing activity. HD5, in contrast, robustly neutralized EV-A71 (50% inhibitory concentration [IC_50_], 8.1 μM; 95% confidence interval [CI], 6.1 to 9.6 µM; Hill slope, -6.5) (**Fig 1B**). However, EV-D111, a member of the EV-D species, was resistant to HD5 neutralization (**Fig 1C**). These findings contrast with prior studies of echovirus 11, a member of EV-B, and demonstrate that EV infection can be sensitive to α-defensin neutralization. However, the anti-EV effects of α-defensins vary depending on the combination of peptide and viral serotype.

**Fig 1:**
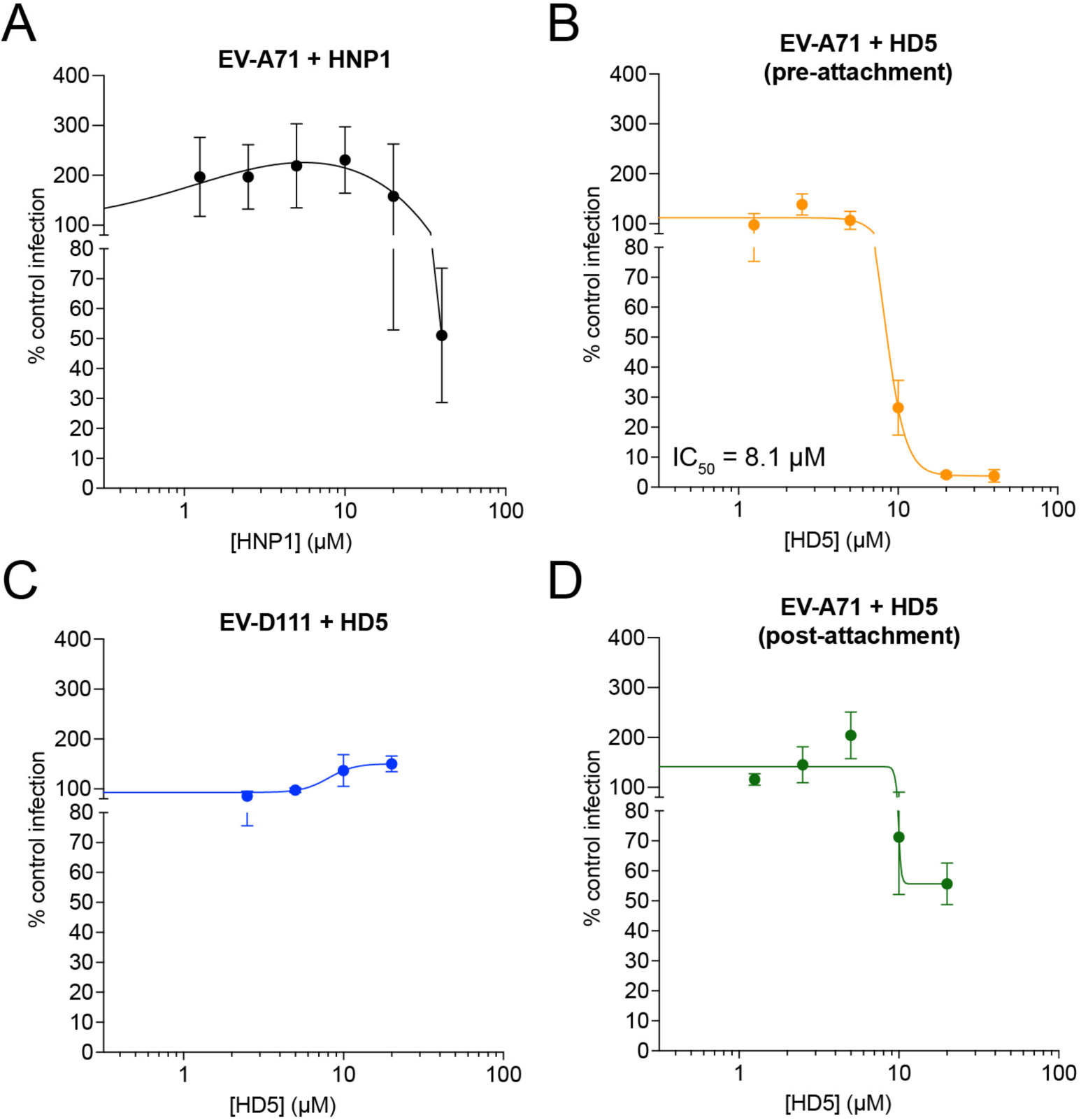
Human α-defensins neutralize EV-A71 but not EV-D111 infection. EV-A71 was exposed to A) HNP1 or B) HD5 on ice prior to infection of RD cells. C) EV-D111 was exposed to HD5 on ice prior to infection of RD cells. D) EV-A71 was exposed to HD5 post-attachment to RD cells. For all assays, data are normalized to control infection in the absence of α-defensin and are the mean ± SD of three to four biological replicates. The mean IC_50_ is indicated in B.

### HD5 disrupts EV-A71 entry

To understand how HD5 disrupts EV-A71 infection, we first performed an order-of-addition assay. Our initial experiments (**Fig 1B**) were conducted under pre-attachment conditions: HD5 was incubated with virus on ice, and the complex was then added to cells. However, if HD5 was added to EV-A71 that was first bound to RD cells in the cold (post-attachment), neutralization was partially attenuated, with only 44% reduction in viral infection at 20 μM HD5 (**Fig 1D**). This suggests that HD5 binds EV-A71 to block infection rather than targeting the host cell.

To determine when in the viral replication cycle EV-A71 is targeted by HD5, we performed a time-of-addition assay. The kinetics of EV-A71 replication in RD cells are well established (**Fig 2A**) (29). In the first 3 h post infection (p.i.) the virus enters the cell, uncoats, and initiates genome translation. Genome replication then begins, followed by assembly and egress between 6 and 24 h p.i. To synchronize infection, we bound EV-A71 to cells in the cold. We then added HD5 while EV-A71 was binding to cells (-1 h p.i.); immediately before initiating infection by warming (0 h p.i.); as well as 1, 3, or 5 h p.i. Note that the -1 h p.i. time point is distinct from pre-attachment conditions (**Fig 1B**), where HD5 is incubated with EV-A71 in the absence of cells, since HD5 and EV-A71 are incubated in the cold in the presence of cells in this assay. Under these conditions, HD5 was only able to neutralize EV-A71 infection when added at -1 or 0 h p.i. (**Fig 2B**). Therefore, HD5 binding to EV-A71 likely blocks entry.

**Fig 2:**
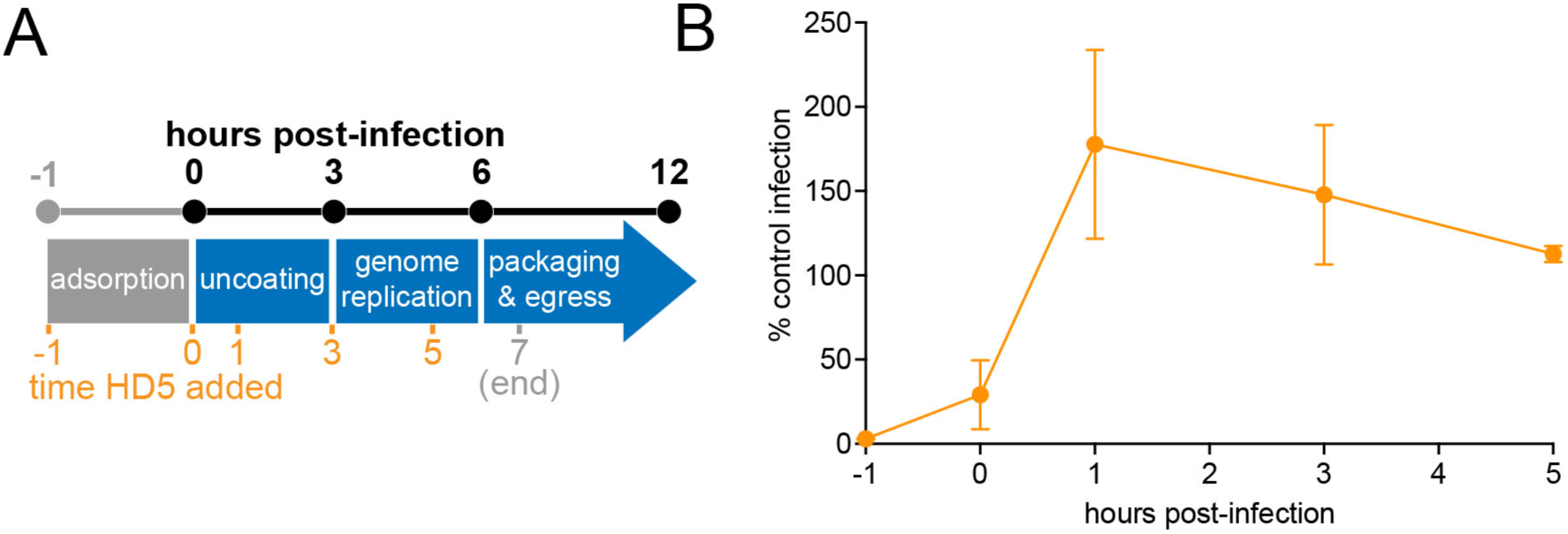
HD5 disrupts EV-A71 entry. A) Schematic of EV-A71 infection kinetics. B) 20 μM HD5 was added at the indicated times (labeled in orange in A). Data are normalized to control infection in the absence of HD5 and are the mean ± SD of three biological replicates.

### HD5 disrupts EV-A71 *in vitro* uncoating and genome release

To confirm HD5 binding to EV-A71, we analyzed the effects of HD5 on uncoating of purified EV-A71 *in vitro* using differential scanning fluorimetry (DSF). The EV-A71 capsid is made up of sixty repeating protomer units, with each protomer consisting of four proteins, VP1 to VP4, and a lipid molecule called “pocket factor” that binds a hydrophobic pocket in VP1 (30–32). In the infectious EV-A71 virion, VP1, VP2, and VP3 are surface exposed, while VP4 is buried within the interior of the capsid. EV-A71 uncoating proceeds via an irreversible series of conformational changes that begins with the externalization of the VP1 N-terminus, loss of VP4 and pocket factor, and expansion of the virion by ∼4% to form the altered “A-particle” uncoating intermediate that still contains genomic RNA (33–36). EV-A71 uncoating can be induced *in vitro* by either incubation with its uncoating receptor, SCARB2, under acidic conditions or by heating and can be monitored using DSF for SYPRO orange fluorescence (35,37).

In good agreement with previous results (35,37), when purified EV-A71 was analyzed by DSF we observed a two-step unfolding process with a melting temperature (T_m_) of 53.5 ± 1.0°C. This corresponds with heat-triggered particle expansion followed by non-specific protein unfolding and aggregation at temperatures greater than 60°C (**Fig 3A and B**) (35). The addition of 20 μM HD5 increased the T_m_ to 55.6 ± 0.5 °C, indicating that HD5 binds directly to the EV-A71 capsid and stabilizes it against heat-triggered particle expansion.

**Fig 3:**
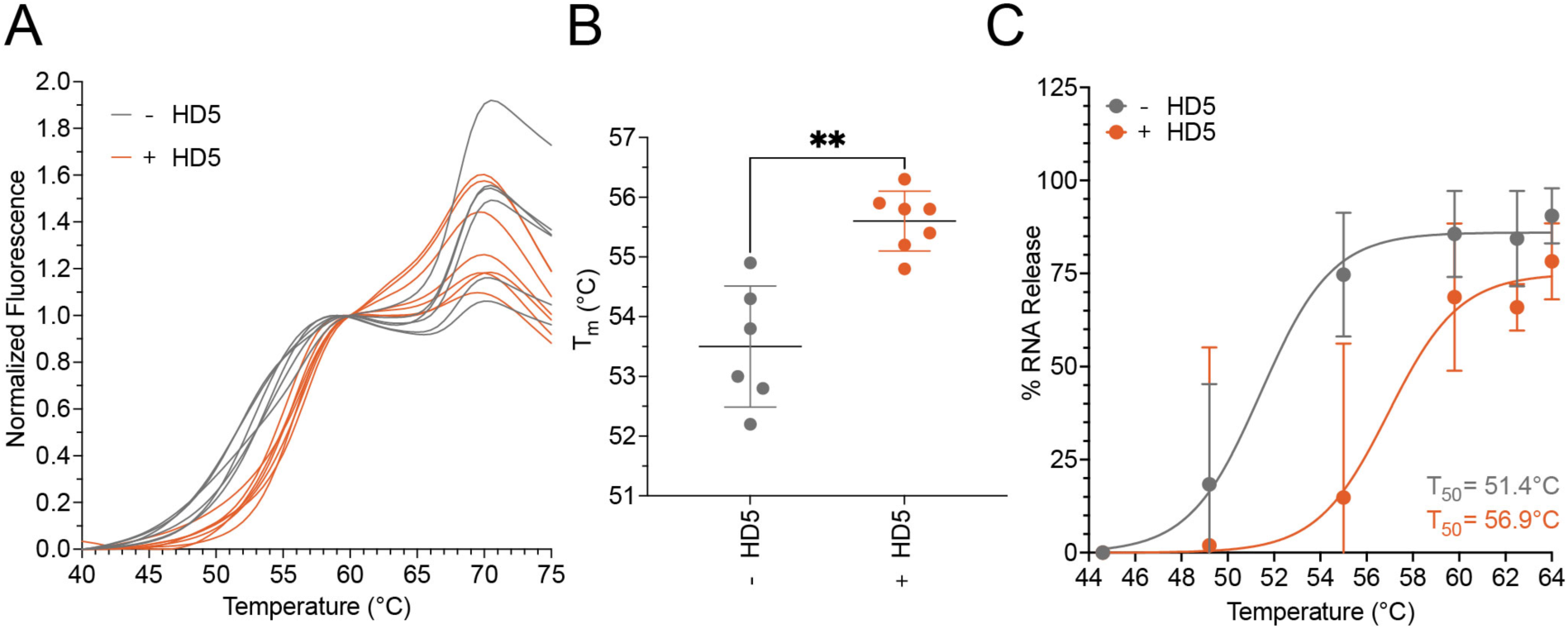
HD5 disrupts EV-A71 *in vitro* uncoating. A) Purified EV-A71 was incubated ± 20 μM HD5, then capsid expansion and unfolding were monitored by DSF. Data are min-max normalized between 40 and 60°C. B) T_m_ values from A. Data are the mean ± SD of six to seven biological replicates. C) RT-qPCR quantification of percent genomic RNA release from unpurified EV-A71 as a function of temperature. Data are normalized to the 44.6°C value for each condition (± 20 μM HD5) then fit to a Boltzmann sigmoidal model. Mean T_50_ values are indicated.

After formation of the A-particle, the RNA genome is externalized, resulting in an empty particle (34,35). To determine if HD5 disruption of viral particle expansion alters RNA genome release, we subjected unpurified EV-A71 that was pre-incubated in the presence or absence of 20 μM HD5 to a temperature gradient, treated samples with Benzonase Nuclease to degrade externalized RNA, and quantified the remaining RNA by RT-qPCR (35). Temperature-dependent RNA release curves were then fitted to the Boltzmann sigmoidal equation to determine T_50_, the temperature at which half of the EV-A71 genome is released. In the presence of HD5 (T_50_, 56.9°C; 95% CI, 54.3-59.8°C), we observed a significant right shift (P < 0.05) compared to EV-A71 alone (T_50_, 51.4°C; 95% CI, 50.0-53.3°C) (**Fig 3C**). This corresponds to a ΔT_50_ of 5.5°C, which is in good agreement with the ΔT_m_ of 2.1°C that we observed in the DSF experiments. Therefore, HD5 binding to the EV-A71 capsid leads to disruption of both particle expansion and genome release.

### Primary, tertiary, and quaternary structure are required for HD5 activity against EV-A71

To define the structural determinants of HD5 anti-EV activity, we tested a series of HD5 analogs for their ability to neutralize EV-A71 infection. Human α-defensins assemble into a predominantly β-sheet tertiary structure stabilized by three intramolecular disulfide bonds, which is required for antiviral activity against other non-enveloped viruses (5,8,9,38–41). To determine if this requirement extends to EV-A71, we used an HD5 analog containing α-aminobutyric acid in place of each cysteine (HD5-Abu). This peptide exhibited no neutralizing activity against EV-A71 (**Fig 4A**). In addition to disulfide bond formation, dimerization is essential to HD5 antiviral activity (8,9). To determine a requirement for HD5 dimerization, we used an obligate monomer HD5 analog in which stabilizing hydrogen bonds at the dimer interface are disrupted by methylation of the peptide bond between C20 and E21 (HD5-E21me) (42). HD5-E21me also exhibited no neutralizing activity against EV-A71 (**Fig 4B**). Together, these studies indicate that a homodimer of the disulfide-stabilized monomer is the active antiviral form of HD5.

**Fig 4:**
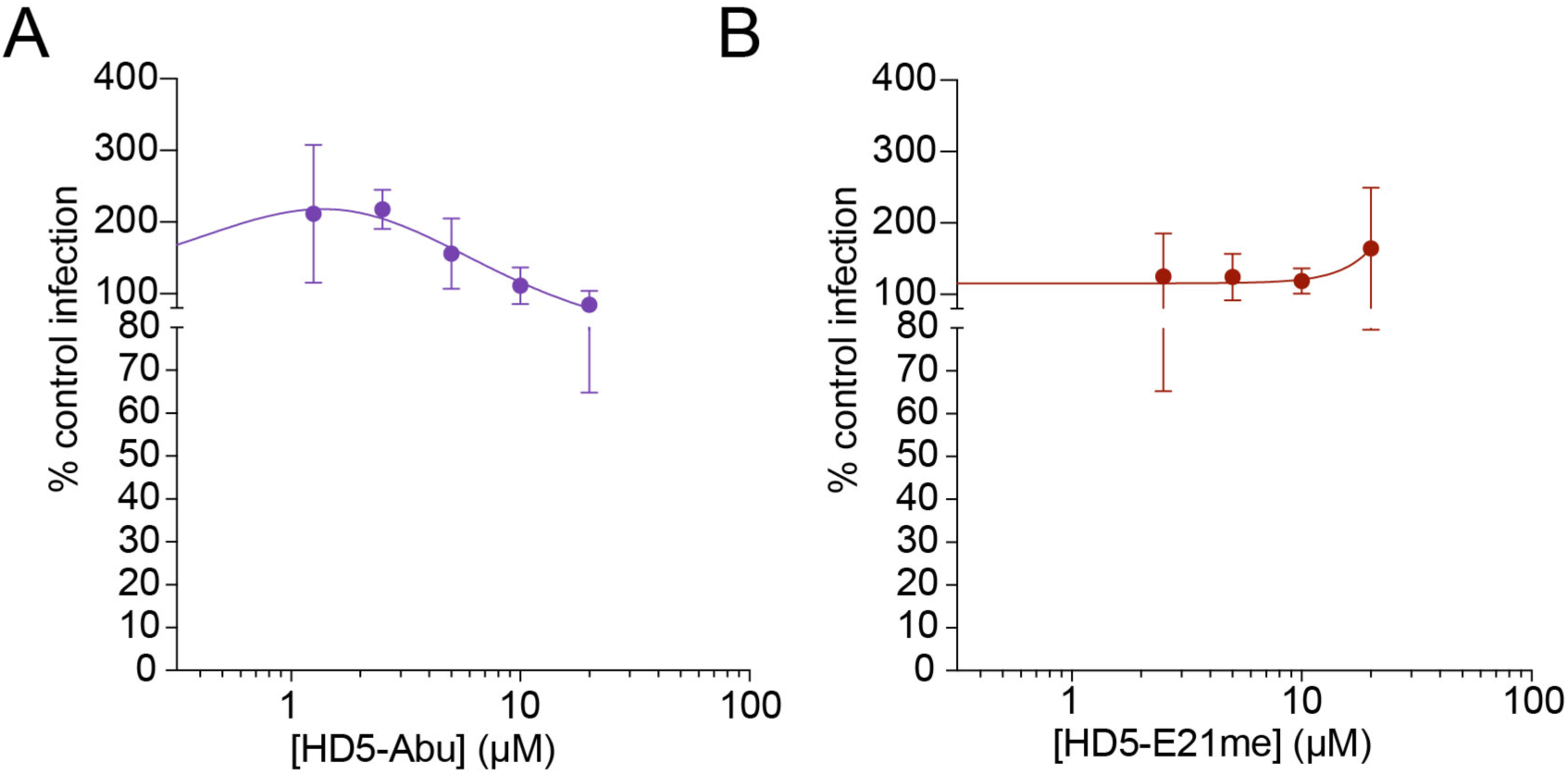
Tertiary and quaternary structure are essential for HD5 activity against EV-A71. A) HD5-Abu lacking disulfide bonds does not neutralize EV-A71 infection. B) The obligate monomer HD5-E21me has no effect on EV-A71 infection. Data are the mean ± SD of three biological replicates.

We next conducted alanine scanning mutagenesis to identify specific HD5 residues that are important for EV-A71 neutralization. We examined single alanine substitution analogs for all HD5 residues except for the six cysteines involved in disulfide bond formation, an invariant glycine (G18) required to form a β-bulge (41,43), a conserved salt bridge (R6 and E14) (44,45), and the two endogenous alanine residues. Eleven of the mutant peptides had no effect on HD5 neutralizing potency, while ten attenuated inhibition (**Fig 5A**): mutation of I22 or R25 led to a > 2-fold increase in IC_50_ (I22A: 11.9 μM, R25: 16.8 μM); mutation of R9, V19 or L26 led to a > 3-fold increase in IC_50_ (R9: 18.4 μM, V19: 17.3 μM, L26: 18.6 μM); while mutation of R13, L16, Y27, R28, or R32 resulted in a complete loss of neutralization compared to wild type HD5 (5.8 μM). We mapped these residues to the structure of the HD5 homodimer. The side chains of R9, L16, V19, R25, L26, R28, along with the main chain carbonyl of Y27 form a single continuous surface (surface A) clustered on one face of the HD5 dimer (**Fig 5B**, left). This surface extends to the opposite face of the dimer and includes the side chains of I22 and Y27 (**Fig 5B**, right). Rotating HD5 by 180° reveals a second smaller surface (surface B) that consists of the R13 and R32 side chains.

**Fig 5:**
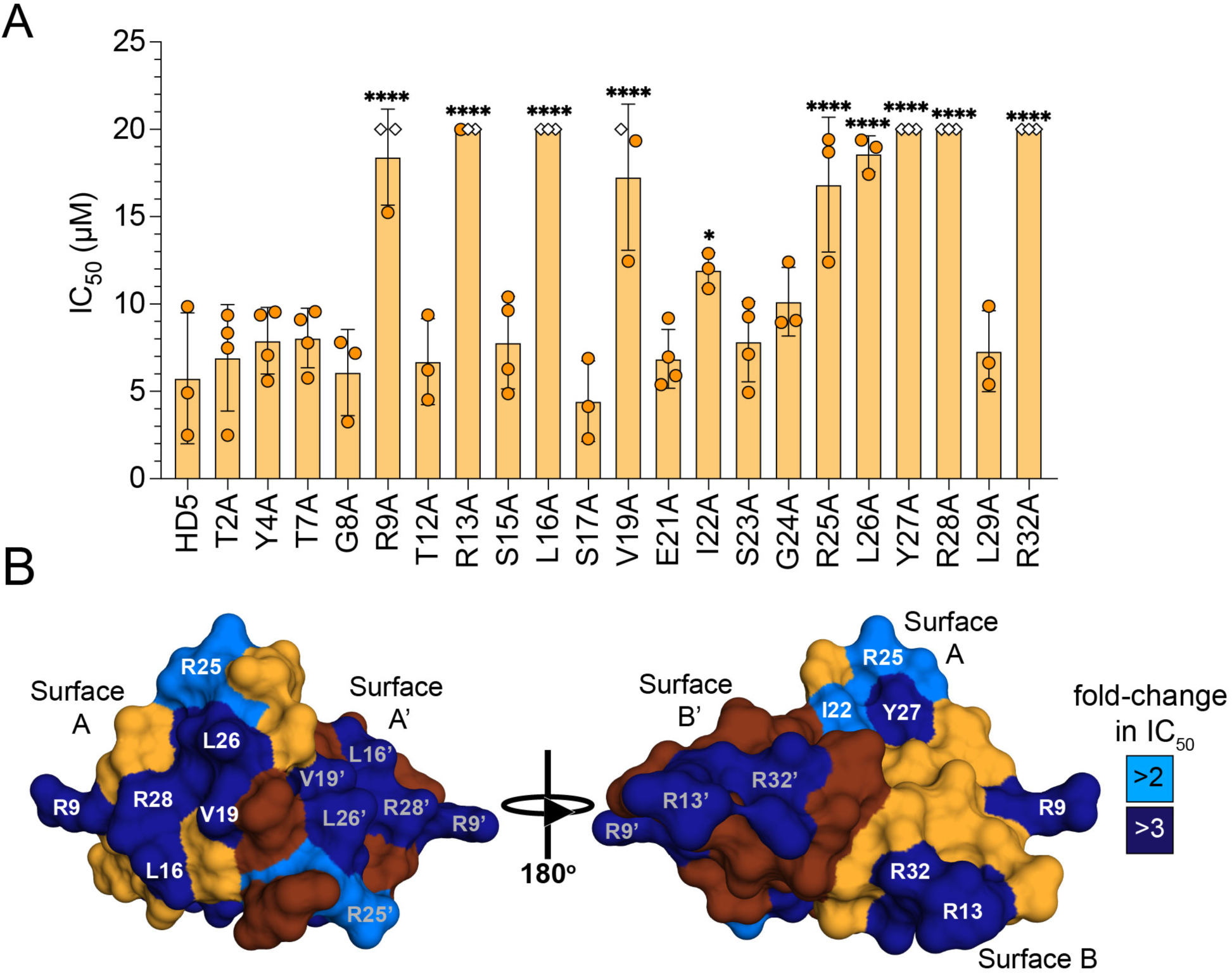
Two surfaces are essential for HD5 anti-enteroviral activity. A) HD5 or single alanine substitution mutants were incubated with EV-A71 on ice prior to infection of RD cells. IC_50_ values were determined separately for each of three to four biological replicates. For replicates with less than 50% inhibition at the highest concentration of peptide tested (white diamond, 20 µM), an IC_50_ of 20 μM was imputed. B) Residues important for antiviral activity are mapped to the HD5 homodimer (PDB: 1ZMP; Chain A orange, Chain B chocolate). Mutations resulting in a > 2-fold shift in IC_50_ are in marine, and a > 3-fold shift are in navy. Surface A (R9, L16, V19, I22, R25, L26, Y27, R28) and B (R13, R32) are labeled on each monomeric chain.

### HD5 disrupts EV-A71 uncoating via two independent pathways

To determine if the HD5 residues important for neutralization of EV-A71 are also involved in capsid binding and disruption of *in vitro* uncoating, we selected seven point mutants (R9A, R13A, V19A, L26A, Y27A, R28A, R32A) that were attenuated >3-fold in antiviral function and assessed their effects on capsid stabilization and genome release. The T_m_ of EV-A71 incubated with each analog fell between that of virus alone (53.5°C) or virus incubated with WT HD5 (55.6°C) (**Fig 6A**). Aside from WT HD5, only R13A significantly raised the T_m_ compared to virus alone (ΔT_m_ = 1.8°C). To measure genome release, we adapted the assay described in **Fig 3C** to monitor RNA release at a single temperature (55.5°C) using purified virus to remove the possibility of confounding contributions of unknown cellular factors. Virus alone released 97.1% of its genome, whereas only 61.8% was released in the presence of 20 μM HD5. Of the mutants tested, both R9A (58.6%) and L26A (50.3%) blocked genome release more efficiently than WT HD5, while R28A (68.2%) and R32A (77.1%) partially blocked genome release. The remaining three peptides (R13A, V19A, Y27A) lost their ability to significantly block genome release. We then compared these results to the infection assay from **Fig 5A** using data obtained at the same peptide concentration (20 µM) across all three assays. At this concentration, HD5 inhibited 91.6% of EV-A71 infectivity. Three peptides (Y27A, R28A, R32A) completely lost inhibitory activity, while the remaining mutants exhibited an intermediate amount of inhibition (**Fig 6C**).

**Fig 6:**
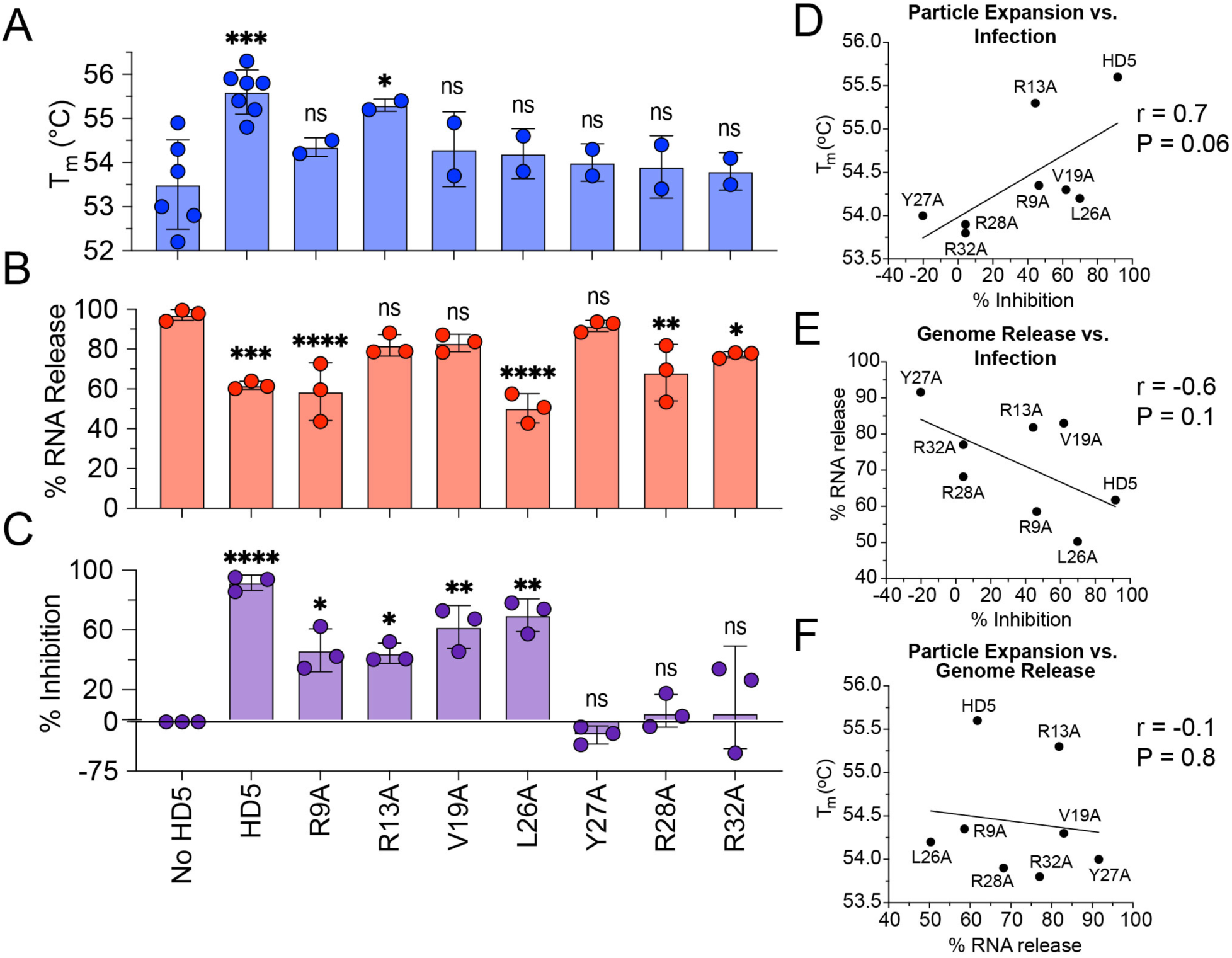
HD5 neutralization of EV-A71 infection is correlated to blocked particle expansion and genome release. A) Purified EV-A71 was incubated ± 20 μM HD5 or HD5 mutants, then capsid expansion and unfolding were monitored by DSF. Data for no HD5 and WT HD5 controls are replotted from Fig 3B. All other data are the mean ± SD of two biological replicates. B) RT-qPCR quantification of percent genomic RNA release from purified EV-A71 ± 20 μM HD5 or HD5 mutants at 55.5°C. Data are the mean ± SD of three biological replicates. C) The data from Fig 5A were used to calculate percent inhibition of unpurified EV-A71 infection upon exposure to 20 μM HD5 or HD5 mutants on ice prior to infection of RD cells. For EV-A71 infection in the absence of HD5, a value of 0% inhibition was imputed for each replicate. Data are the mean ± SD of three biological replicates. The mean values from A to C were plotted against each other to compare inhibition of D) infection and particle expansion (T_m_), E) infection and genome release (percent RNA release), and F) particle expansion and genome release. Pearson’s r values and P values are indicated on each plot.

Upon analyzing each peptide across all three assays, several distinct phenotypes emerged. Both WT HD5 and the R13A mutant stabilized the capsid and inhibited infection, suggesting that capsid stabilization may be sufficient for neutralization. To test this across all peptides, we replotted the data from **Fig 6A** and **Fig 6C** into a correlation plot and calculated Pearson’s r (46). There is a moderate positive correlation between T_m_ and % inhibition (r = 0.7, R^2^ = 0.5, P = 0.06, 95% CI -0.04-0.9), supporting a mechanism where capsid stabilization leads to neutralization (**Fig 6D**). In contrast, the correlation between genome release and inhibition of infection is more variable. Of the four peptides that inhibited genome release to some degree, two (R9, L26) blocked infection and two (R28, R32) did not. Nevertheless, there is a moderate negative correlation between genome release and inhibition (r = -0.6, R^2^ = 0.3, P = 0.1, 95% CI - 0.9-0.2) (**Fig 6E**). These data suggest that inhibiting particle expansion or genome release represents two independent pathways to disrupting EV-A71 uncoating. Consistent with this hypothesis, T_m_ and genome release are poorly correlated (r = -0.1, R^2^ = 0.02, P = 0.8, 95% CI -0.8-0.6) (**Fig 6F**). We therefore propose that the block to EV-A71 uncoating can proceed via two parallel mechanisms, both of which are necessary to fully neutralize EV-A71 infection (**Fig 7**).

**Fig 7:**
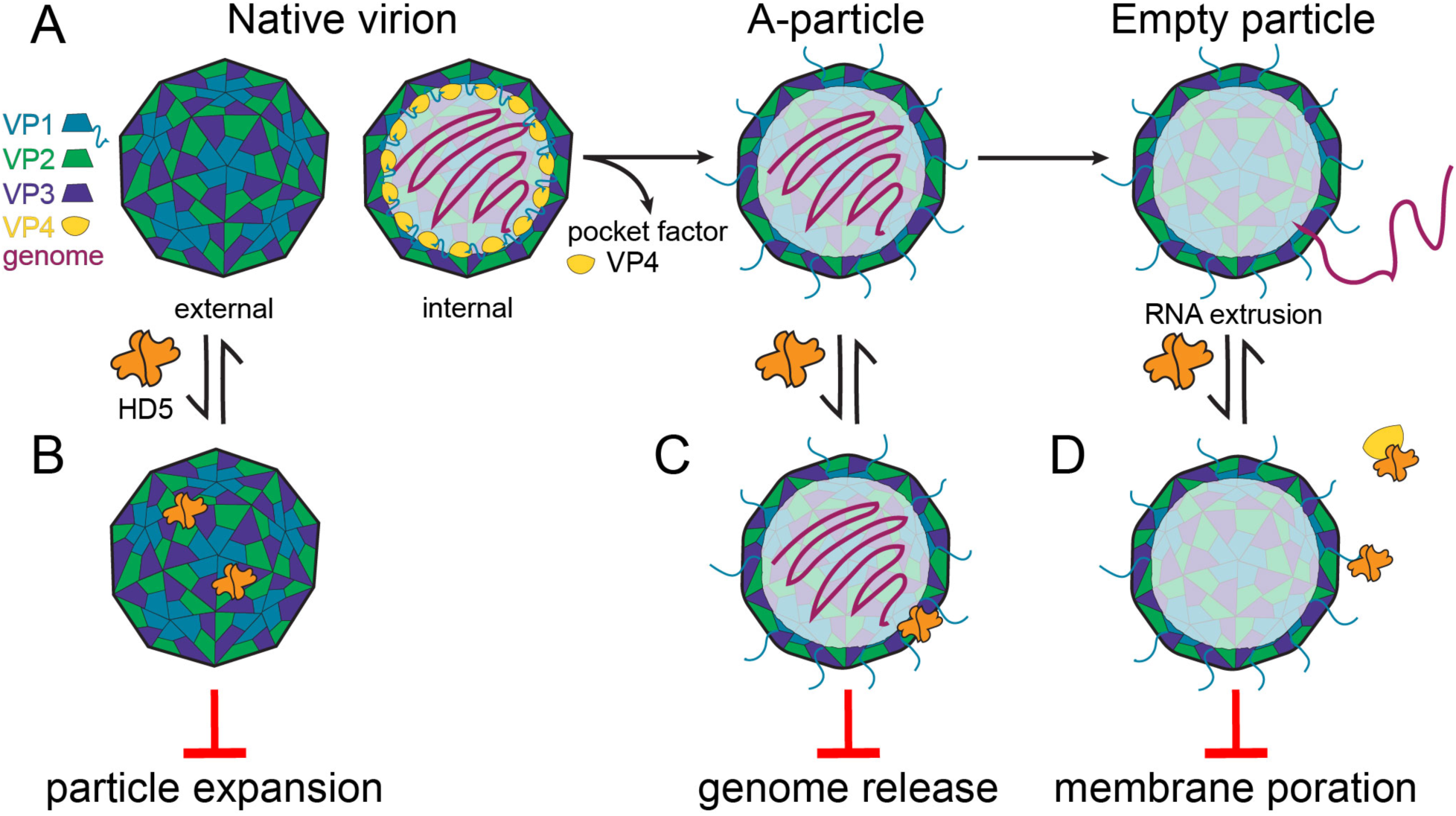
Mechanism of HD5 action against EV-A71. A) EV-A71 uncoating begins with expulsion of pocket factor and VP4, leading to conformational changes that result in externalization of the VP1 N-terminus and formation of the expanded A-particle. RNA is then extruded through a negatively charged channel, yielding the empty particle. B) Proposed binding mode 1: HD5 may bind to the native virion and disrupt pocket factor release and protomer rotation, leading to blocked particle expansion. C) Proposed binding mode 2: HD5 may bind to the negatively charged channel and block genome release. D) Proposed binding mode 3: HD5 may bind externalized VP1 N-termini and/or VP4 and block membrane poration and RNA release into the host cytosol.

## Discussion

We determined that HD5 neutralizes EV-A71 infection, providing the first example of an EV that is sensitive to α-defensins and expanding the scope of α-defensin antiviral activity to a new family of pathogenic RNA viruses. We propose HD5 disruption of EV-A71 uncoating as a mechanism to explain neutralized infection in cells, since HD5 binds to the EV-A71 capsid and stabilizes the virus against heat-triggered uncoating, blocking particle expansion (ΔΤ_m_ = 2.1°C) and genome release (ΔT_50_ = 5.5°C). It is generally accepted that heat-triggered conformational changes in EV-A71 faithfully recapitulate viral cellular uncoating (34,35,47–49). Stabilization of the EV-A71 capsid via mutagenesis or inhibitor binding typically leads to *in vitro* thermal shifts ranging from 2 to 8°C, in good agreement with our observed values, and these perturbations lead to disrupted infection in cells (35,50–54). Therefore, a well-established relationship exists between increased *in vitro* EV thermostability and disrupted uncoating, suggesting that our *in vitro* observations expand to a cellular mechanism. This notion is supported by the fact that disruption of both particle expansion and genome release correlates with inhibition of infection for HD5 and HD5 mutants. Future studies interrogating EV-A71 cellular entry and uncoating in the presence of HD5 would provide additional insights into this mechanism.

We utilized a suite of HD5 mutants to dissect the structural features important for activity against EV-A71, expanding upon previous studies with human adenovirus 5 (HAdV-5) and human papillomavirus 16 (HPV16) and revealing several conserved elements broadly involved in HD5 antiviral activity (8,10). Biochemical studies confirm that the HD5 mutants utilized in these studies are properly folded, ruling out global structural abnormalities as an explanation for attenuated anti-EV activity (42). As has been true for all viruses that have been tested including HAdV, HPV, adeno-associated virus (AAV), and SARS CoV-2, disulfide-stabilized tertiary structure is essential for activity against EV-A71 (7,9,10,12,14,15,55). Moreover, a requirement for homodimerization is supported by the lack of activity of both HD5-E21me and HD5-Y27A, both of which are monomeric (8–10,56). Beyond tertiary and quaternary structure, α-defensins are both cationic and amphipathic, and their antimicrobial activity is mediated by electrostatic and hydrophobic interactions with their targets (8,42,57–61). Mutating any of the HD5 arginines to alanine leads to at least a > 2-fold shift in IC_50_ against of EV-A71, confirming a role for positive charge in HD5 anti-EV activity. However, mutating the sole acidic residue (E21) to alanine does not increase HD5 potency. Moreover, HNP1 (+3) is only slightly less positive than HD5 (+4) yet fails to neutralize EV-A71. Thus, there is no simple relationship between charge and activity. In the case of HAdV-5 and HPV16, hydrophobicity is also an important contributor to antiviral potency (8,10). Similarly, several hydrophobic residues including L16, V19, I22 and L26 are important for HD5 anti-EV activity. Thus, like other viruses, HD5 anti-EV activity is mediated by a complex interplay between primary sequence, tertiary structure, and quaternary oligomerization. Some of these features may also explain why HD5 but not HNP1 neutralizes EV-A71.

The ten HD5 residues important for anti-EV activity cluster to two continuous surfaces: a large “surface A” that extends across one face of the HD5 dimer and a smaller “surface B” located on the opposite face. Every residue within surface A except for R9 is also important for HD5 neutralization of HAdV-5 and HPV16, suggesting this may be a conserved interface involved in broad-spectrum HD5 antiviral activity (10). None of the surface A mutants tested in DSF experiments were able to block particle expansion, suggesting HD5 may bind directly to the EV-A71 capsid via this interface and that the integrity of this interface in its entirety is required for this function. The effects of mutations within surface B are more variable: R13 is dispensable for capsid stabilization while R32 is required. Therefore, surface B may also be involved in direct binding to the EV-A71 capsid, despite being located on the opposite face of the HD5 dimer. HD5 binds to both HAdV and AAV at very high stoichiometry, with hundreds if not thousands of copies of HD5 bound to each virion, and variability in antiviral potency between HD5 mutants is driven by changes to capsid binding stoichiometry (7,9,10,12,13). This high stoichiometry also suggests a role for higher order oligomerization of HD5 as a requirement for antiviral activity. HD5 may also bind to EV-A71 at high stoichiometry and access multiple binding modes, both to the capsid and to other HD5 molecules, which could explain how surface A and B could both be involved in capsid stabilization. While surface A may be a conserved interface broadly important for antiviral activity, there are undoubtedly differences in how HD5 interacts with each specific virus. This may explain why several residues (T2, Y4, T7, E21, G24, L29) are required for HD5 activity against HAdV-5 and HPV16 but not EV-A71 (10). Nevertheless, the presence of a conserved interface that is adaptable to the surface topology of disparate viral capsids through multimodal binding may explain why HD5 can achieve broad-spectrum antiviral activity.

Our data suggest that HD5 has two functions, disruption of EV-A71 capsid expansion and genome release, that are separable and both required for complete neutralization. EV-A71 uncoating begins with the expulsion of pocket factor and conformational changes within VP1, resulting in rotation of the capsid protomers around each other and opening of a small channel through which the VP1 N-terminus externalizes (**Fig 7A**) (31,33,34). A large negatively charged channel also opens nearby, which is thought to allow VP4 release and downstream RNA extrusion. HD5 binding could block pocket factor expulsion or protomer rotation, which would disrupt both particle expansion and genome release (**Fig 7B**). A block in genome release without capsid stabilization (as observed for R9A and L26A) could be explained by an interaction between HD5 and the negatively charged interior of the RNA channel to sterically block genome extrusion, or by interactions with the RNA itself (**Fig 7C**). In DSF experiments, the externalized hydrophobic VP1 N-terminus or VP4 regions may contribute to SYPRO orange fluorescence. Therefore, one possible explanation for capsid stabilization without disrupted genome release (as observed for R13A) is that HD5 binds to VP1 and/or VP4 after viral uncoating, blocking SYPRO orange binding and raising the apparent T_m_ (**Fig 7D**). Both the VP1 N-terminus and VP4 are thought to be essential for poration of the endosomal membrane to release the RNA genome into the cytosol, so blocking these elements would still lead to disrupted EV-A71 entry (62–64). The EV capsid is dynamic, and the native virion samples reversible conformations in which internal capsid elements are transiently exposed (19,65). Therefore, HD5 may access cryptic capsid binding sites either by exploiting this conformational equilibrium or by accessing new binding sites that are exposed in the endosome as the virus begins to uncoat. Because HD5-induced capsid stabilization and blocked genome release are independent and both phenotypes are required for full neutralization of infection, it is likely HD5 accesses several of these binding sites.

EV capsid stabilization and disrupted uncoating are common antiviral inhibitor mechanisms. For example, small molecules that compete for pocket factor binding to VP1 stabilize the EV capsid and disrupt both *in vitro* and cellular uncoating (50–52,54,66–76). Furthermore, peptides derived from cathelicidins, a family of antimicrobial peptides distinct from defensins, bind to the capsid exterior and block virus-receptor interactions, thereby disrupting uncoating and infection (53). Therefore, HD5 shares an overall conserved mode of action with these inhibitors. However, to our knowledge the R9A and L26A mutants are unique examples of inhibitors that allow for particle expansion but still block downstream genome release. Our proposed R13A binding mode (**Fig 7D**) is a distinct antiviral mechanism compared to known anti-EV agents. These HD5 mutants therefore offer an opportunity to develop new methods for targeting EV uncoating.

The mechanism of EV-A71 neutralization that we propose shares several parallels with what has been previously observed for DNA viruses. For HAdV, AAV, and HPV, α-defensins bind directly to the viral capsid and inhibit entry by disrupting capsid or genome trafficking through the cell (7–9,11–16,77,78). In the case if HAdV and AAV, HD5 binding blocks key conformational changes associated with uncoating and/or endosomal escape, while for HPV16, HD5 blocks genome release from the endosome and redirects it to the lysosome. HD5 binding to JC polyomavirus (JCPyV) also results in disrupted capsid conformational change and blocked trafficking to the endoplasmic reticulum, thus following the same general mechanism as observed for HAdV, AAV, and HPV (17). The cellular antiviral mechanisms of α-defensins, therefore, represent variations on a common theme of disrupted viral uncoating and/or trafficking that is absolutely dependent on HD5 binding to the viral capsid. EV-A71 neutralization also involves HD5 binding to the capsid and disruption of viral entry, suggesting that some features of the antiviral mechanisms described for DNA viruses extend to RNA viruses. However, unlike DNA viruses, EV-A71 does not traffic through the cell but rather releases its genome directly into the cytosol from the endosome (19). Though further validation is required to confirm our proposed mechanism in cells, this work suggests that capsid stabilization can still lead to disrupted infection of EV-A71 by blocking genome release into the cytosol despite the distinct entry pathways utilized by DNA and RNA viruses. While genome translocation from the endosome is currently the prevailing model for EV uncoating, an alternative model has been suggested involving virion escape into the cytosol prior to genome release (36). HD5 and HD5 mutants could therefore serve as tools to further elucidate the mechanism of EV uncoating. Overall, this work has uncovered several common features of HD5 antiviral action across viral families and advances our understanding of how α-defensins interact with an important enteric pathogen.

## Materials and methods

### Cells and viruses

Human rhabdomyosarcoma (RD) cells were a kind gift from Michael Gale Jr. of the University of Washington (Seattle, WA). Cells were maintained in Dulbecco’s modified Eagle medium (DMEM) supplemented with 10% Fetal Bovine Serum (FBS), 4 mM L-glutamine, 0.1 mM nonessential amino acids, 100 units/mL penicillin and 100 μg/mL streptomycin (complete DMEM). Enterovirus 71 (EV-A71) strain USA/2018-20932 (Cat: NR-52000; Lot: 70032015) and EV-D111 strain ANG/2010/23294 (Cat: NR-52318; Lot: 70043450) were obtained from BEI Resources. Viral working stocks were prepared in RD cells in supplemented DMEM lacking FBS (Serum Free Media, SFM) at an MOI of approximately 1 pfu/cell. When complete cytopathic effect (CPE) was observed (approximately 1-3 d p.i.), cells and media were collected, disrupted by three cycles of freeze/thaw lysis, centrifuged for 10 min at 25,000 × *g* to remove cell debris, and flash frozen. P2-P4 viral stocks were used for all experiments.

### Viral purification

EV-A71 was purified as previously described with the following modifications (79). RD cells were infected at an MOI of approximately 0.1 pfu/cell in supplemented DMEM containing 2.5% FBS. When complete CPE was observed (approximately 48 h p.i.), cells and media were pooled into 500 mL centrifuge tubes and disrupted by three cycles of freeze/thaw lysis. The lysate was centrifuged at 4°C for 15 min at 16,300 × *g* using a JA-14 rotor (Beckman Coulter, Inc.). Virus was precipitated for 16-18 h on ice from the cleared supernatant using 8% PEG-8000 and 0.5 M sodium chloride followed by centrifugation at 4°C for 1 h at ∼3,800 × *g* using a Beckman SX4750A rotor. The supernatant was discarded, and the pellet was resuspended in 15 mL purification buffer (10 mM Tris-HCl, 200 mM NaCl, 50 mM MgCl_2_, pH 7.5). 75 μL of 10 mg/mL DNase I was added, the solution was incubated at RT for 10 min, and the reaction was halted by the addition of 4 mL 0.5 M EDTA. The solution was neutralized with ammonium hydroxide and centrifuged at 4°C for 5 min at ∼3,800 × *g* using a Beckman SX4750A rotor. The supernatant was distributed into centrifuge tubes and underlaid with a 1 mL cushion per tube of 30% sucrose (w/v) in purification buffer. Samples were centrifuged at 4°C for 2 h at 288,000 × *g* using a Beckman SW-41ti rotor. Pelleted virus was resuspended and combined to a final volume of 1 mL in purification buffer, layered onto a step gradient consisting of 10-35% potassium tartrate (w/v) in purification buffer in 5% increments, and centrifuged at 4°C for 2 h at 221,000 × *g* using a Beckman SW-41 rotor. Fractions were collected from the bottom of the gradient via puncture, buffer exchanged and concentrated into purification buffer using a 50k MWCO Amicon spin filter, separated by SDS-PAGE, and stained with AzureRed Fluorescent Total Protein Stain. Fractions containing mature virions without contaminating procapsids were pooled, and total protein concentration was quantified using a Bio-Rad Protein Assay. Purified virus was snap frozen in liquid nitrogen, stored at -80°C, and used where specified.

### α-defensin peptides

HD5 and HNP1 were purchased as linear peptides from CPC Scientific, subjected to oxidative refolding, and purified by RP-HPLC as previously described (18). Purity was assessed by analytical RP-HPLC and mass spectrometry. Final peptide stocks were produced in sterilized water and quantified by UV absorbance at 280 nm. Synthesis, purification and validation of HD5-Abu, HD5-E21me, and HD5 alanine mutants have been previously described (42,80).

### Viral neutralization assays

We employed two protocols, which differed in the order of addition of HD5 to virus relative to virus binding to RD cell monolayers in clear bottom, black wall 96-well plates. For all protocols, virus was used at a concentration previously determined to yield 50% (EV-D111) or 90% (EV-A71) maximal signal. For the pre-attachment protocol, virus was incubated with a dilution series of α-defensin in SFM on ice for 45 min. RD monolayers were washed three times with SFM, the virus/defensin mixture was added to cells, and the plate was incubated at 37°C. For the post-attachment protocol, cells were washed on ice with cold SFM, and virus diluted in SFM was added. After incubation at 4°C for 45 min, cells were washed twice with cold SFM, and serially diluted α-defensin was added. Plates were incubated for 45 min at 4°C and then shifted to 37°C. For both protocols, after 1 h incubation at 37°C, virus/defensin was removed, cells were washed twice with complete DMEM, the media was replaced with complete DMEM, and samples were incubated at 37°C. For time of addition assays, monolayers were washed twice on ice with cold SFM then EV-A71 was added along with 20 μM HD5 at indicated time points. **-1 h p.i.:** before adsorption, HD5 was mixed with EV-A71 and the mixture was added to cells and incubated for 1 h at 4°C, washed twice with SFM, then maintained in 20 μM HD5. **0 h p.i.:** After viral adsorption and subsequent washing with SFM, 20 μM HD5 was added to cells immediately before warming to 37°C. **1, 3, 5 h p.i.:** At each timepoint, SFM was replaced with 20 μM HD5 then infection was allowed to continue. For all assays, at 7 h (EV-A71) or 19 h (EV-D111) p.i. cells were fixed with 2% paraformaldehyde in PBS at RT for 20 min, washed three times with PBS, then quenched and permeabilized with 20 mM glycine and 0.5% Triton-X 100 in PBS at RT for 20 min. Cells were stained at RT for 1 h with either the Light Diagnostics Pan-Enterovirus Reagent (1:40 dilution, Cat: 3360, MilliporeSigma) or the anti-VP3 monoclonal antibody L66J (250 ng/mL, Cat: MA518206, ThermoFisher Scientific) in PBS containing 0.05% Tween 80 and 1% w/v bovine serum albumin. Cells were washed three times with PBS containing 0.05% Tween 80 (PBS-T) then stained at RT for 45 min with Alexa Fluor 488-conjugated goat anti-mouse antibody (2 μg/mL, Cat: A-11001, ThermoFisher Scientific). Cells were washed once with PBS-T and twice with PBS, and monolayer fluorescence was quantified using a Sapphire (Azure) variable-mode imager. Background-subtracted total well fluorescence was quantified using Fiji (ImageJ) version 1.53a, and wells were normalized as percent infection compared to controls infected in the absence of α-defensin.

### Differential Scanning Fluorimetry (DSF)

This assay was described previously, with the following modifications (37). Samples (47 μL) consisting of 4-12 ng/μL purified EV-A71 and 20 µM defensin (HD5 or HD5 analogs) in PBS supplemented with 2 mM MgCl_2_ in white low-profile PCR strip tubes (Bio-Rad Cat:TLS0851) with optical caps (BioRad Cat: TCS0803) were incubated on ice. After 45 min, 3 μL of 1% SYPRO orange protein stain (MilliporeSigma Cat: S5692) was added to each sample. After incubation for 15 min on ice, samples were heated from 4°C to 99°C, ramping at 0.5°C per 10 s step, using a Bio Rad CFX96 Real-Time PCR system.

### RNA Release Assay

Samples of EV-A71 in 100 μL PBS supplemented with 2 mM MgCl_2_ were incubated for 45 min on ice in the presence or absence of 20 μM HD5. Individual samples were then incubated for 1 h at 44.6°C, 49.2°C, 55.0°C, 59.8°C, 62.5°C or 64.0°C (**Fig 3C**) or 55.5°C or on ice (**Fig 6B**) using a BioRad C1000 Thermal Cycler, cooled on ice for 5 min, and treated with 1 unit Benzonase Nuclease (MilliporeSigma Cat: 70664). After incubation for 30 min at 37°C, Benzonase was inactivated with 5 mM EDTA. Viral RNA was extracted using the Quick-RNA Viral Kit (Zymo Research Cat: R1035), eluted in 20 μL nuclease free H_2_O, and converted to single-stranded cDNA using the GoScript^TM^Reverse Transcriptase kit (Promega Cat: A5001) with a VP1-reverse primer (5’-AAGAGTGGTGATCGCCGTG-3’). qPCR was performed on a BioRad CFX96 Real-Time PCR system (amplification sequence: 95°C for 3 min followed by 39 cycles of 95°C for 10 s, 55°C for 30 s) using 20 µL samples containing 5 μL cDNA, 1x SsoAdvanced Universal SYBR Green Supermix (BioRad Cat:1725274), and 0.4 μM each forward (5’-GAGTAGAGCTCTCACTCAAGCTC-3’) and reverse (5’-AACTATCGAGAGTGGTCTCAGCT-3’) primer. Primers were designed to amplify the VP1 gene of GenBank MK652139. Note that unpurified EV-A71 was used in **Fig 3C** at a dilution chosen to yield a C_t_ value between 18 and 20 for the 44.6°C sample, while purified EV-A71 was used at 0.13 ng/μL for **Fig 6B**. Data were exported as C_t_ values and normalized to the 44.6°C sample (thermal gradient) or ice controls (analysis at 55.5°C) by calculating ΔC_t_ = C_t_ – C_t(control)_ then calculating % RNA Release = 100 - (100 x (2^-ΔCt^)).

### Statistical analysis

All statistical analysis was performed with GraphPad Prism 10.5.0. For **Fig 1B**, **1C**, **1D**, **4B**, and **5A** IC_50_ values were determined by fitting a four-parameter dose-response curve with variable slope. For **Fig 1B**, the hill slope from four parameter fitting (-6.5) was then fixed to obtain 95% CI values. For **Fig 5A** curves were fit separately for each biological replicate with bottom constraint of zero, then IC_50_ values were plotted together as bar graphs. **Fig 1A** and **4A** were fit using a Bell-shaped curve as a guide to the eye, but these were not used for quantification. For **Fig 3A**, **3B**, and **6A** baseline subtraction of dye-only or defensin-only controls from the raw RFU vs. temperature data was performed then data between 40°C and 75°C were fit to Model 3 and 4 using DSFworld (81). Fit curves were manually inspected to select the best Model for each data set, and the resultant T_m_ values were plotted together. **Fig 3C** was fit to a Boltzmann sigmoidal model with bottom constraint of zero and constant slope of 1.5. **Fig 6D to F** were fit to a simple linear regression as a guide to the eye but not used for quantification. Statistical significance of **Fig 3B** was assessed using an unpaired two-tailed Welch’s t-test. **Fig 3C** was analyzed using the extra sum-of-squares F-Test (P<0.05). Pearson correlation was determined for **Fig 6D to F**. **Fig 5A**, **6A**, **6B**, and **6C** was analyzed using ordinary one-way ANOVA with multiple comparisons to WT HD5 (**Fig 5A**) or the no HD5 condition (**Fig 6A to C**). For all graphs, significance is marked by asterisks: *P<0.05, **P<0.005, ***P<0.0005, ****P<0.0001.

## Acknowledgements

The authors thank Wuyuan Lu (Institute of Human Virology, University of Maryland School of Medicine, Baltimore, MD) for providing the HD5 alanine scan library used in these studies. This work was supported by R01 AI104920 (to J.G.S.) and F32 AI178920 (to K.R.H.) from the National Institute of Allergy and Infectious Diseases, T32 HD007233 (to K.R.H.) from the Eunice Kennedy Shriver National Institute of Child Health and Human Development, and by the National Institutes of Health Office of the Director under Award Number S10 OD026741 (to J.G.S.). The content is solely the responsibility of the authors and does not represent the official views of the National Institutes of Health.

## Notes

### Competing Interest Statement

The authors have declared no competing interest.

